# Empirical lognormality of biological variation: implications for the ‘zero-force evolutionary law’

**DOI:** 10.1101/2020.04.28.066563

**Authors:** Philip D. Gingerich

## Abstract

The zero-force evolutionary law (ZFEL) of McShea et al. states that independently evolving entities, with no forces or constraints acting on them, will tend to accumulate differences and therefore diverge from each other. McShea et al. quantified the law by assuming normality on an additive arithmetic scale and reflecting negative differences as absolute values, systematically augmenting perceived divergence. The appropriate analytical framework is not additive but proportional, where logarithmic transformation is required to achieve normality. Logarithms and logarithmic differences can be negative but the proportions they represent cannot be negative. Reformulation of ZFEL in a proportional or geometric reference frame indicates that when entities evolve randomly and independently, differences smaller than any initial difference are balanced by differences larger than the initial difference. Total variance increases with each step of a random walk, but there is no statistical expectation of divergence between random-walk lineages.

## 1. Introduction

The idea of a zero-force evolutionary law or ‘ZFEL’ was first proposed by McShea (2005) [1] and Brandon (2006) [2], and then developed collaboratively in two co-authored books (McShea and Brandon 2010; Brandon and McShea 2019) [3, 4]. Quantitative formulation of the zero-force law was published recently (McShea et al., 2019) [5]. The basic principle is that “Given two independently evolving entities, in the absence of forces or constraints acting on the differences between them, they will tend to accumulate differences and therefore to diverge, that is, to become ever more different from each other… The ZFEL is a null model. It describes what happens in evolutionary lineages when forces and constraints are absent.” (McShea et al., 2019, p. 1101) [5].

McShea et al. (2019)[5] wrote further “Some may find the result … puzzling, simply because when change is random, the [expectation] for each lineage does not change over time. And it might seem that if the expected values for … two entities do not change, so that the distance between their expected values is constant, then the expected (absolute) distance between the entities … should not change. In fact it does. It increases.” (McShea et al., 2019, p. 1103)[5].

Quantitative formulation of the zero-force law is important for making the law both operational and testable. McShea et al. (2019)[5] stated that diversity and complexity are variance measures, and that variances increase as random events accumulate (which is true), but the model they developed to quantify ZFEL is not about variances. Their model quantifies an expected or mean difference between random-walk lineages as they evolve in parallel through time.

Difference can refer to *arithmetic difference,* the signed subtraction difference between two amounts or values on a number line, e.g., *a – b* or *b – a*, and it can refer to *absolute arithmetic difference* or distance, the positive absolute value of an arithmetic difference, e.g., |*a – b*| = |*b – a*|. Absolute difference is the measure employed by McShea et al. (2019)[5].

Alternatively, and importantly, difference can refer to *proportional difference,* the ratio of two amounts, e.g., *a* / *b* or *b* / *a*. A proportional scale is a logarithmic scale, where ratios are represented as differences: ln (*a* / *b*) = ln *a* – ln *b*, and ln (b / a) = ln *b* – ln *a*. Measures of the entities involved, populations of organisms or species, cannot be negative or zero. On a proportional scale differences can range from infinitesimal when they are very small, to 1 when the entities are identical (*a* / *a* = 1), to infinite when differences are very large — but there are no negative differences of proportion.

McShea et al. (2019)[5] formulated the zero-force evolutionary law in terms of absolute difference between lineages analyzed on an arithmetic scale. They acknowledged (p. 1107) that “in this model, change is additive, but because change in biology is often proportional, the horizontal axis could alternatively be interpreted as a log scale.” However, McShea et al. (2019)[5] did not pursue this alternative. If they had pursued it they would have discovered that their zero-force evolutionary law cannot be justified on a proportional scale. A zero-force law based on proportional difference is developed here, first to promote a better understanding of the ubiquity of proportional difference, and second to illustrate the essential role logarithms play in evolutionary analysis.

## 2. Arithmetic lognormality and geometric normality

*Loi des erreiirs… tout le monde y croit … les expérimentateurs s’imaginent que c’est un théorème de mathématiques, et les mathématiciens que c’est un fait expérimental.* [Law of errors. everyone believes it … empiricists think it is proven mathematically, and mathematicians think it is demonstrated empirically.] Henri Poincaré, *Calcul des Probabilités,* 1896, p. 149.[6]

Natural selection acts on variation in populations of organisms. The variation can be quantified by counting or measuring characteristics like limb length or body weight in a sample from a population. Empirical measurements have a central value and a distribution of variation about this. The sample mean, 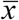, and the sample standard deviation, *s*, describe the location and dispersion of a corresponding normal curve, the biological manifestation of Henri Poincaré’s ‘law of errors.’ When biological measurements represent areas or volumes and samples span a range of sizes it becomes clear that *s* increases with 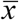. Biometrician Karl Pearson (1896)[7] recognized this and proposed a ‘coefficient of variation,’ 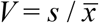, to standardize comparisons of dispersion across samples.

Standardizing dispersion is not the only problem. Most studies of phenotypic variation compare linear measurements in similar populations, where measurements yield bell-shaped distributions and any small error of modeling the distributions as normal is ignored because it is difficult to see (figure 1a). When the coefficient of variation *V* is larger and populations are well sampled, empirical measurements are positively skewed (figure 1c-f). Correlation of *s* with 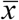, and skewness at high *V* expose deficiencies in the regularity and symmetry expected for simple normality on an arithmetic scale of measurement.

**Figure 1.**
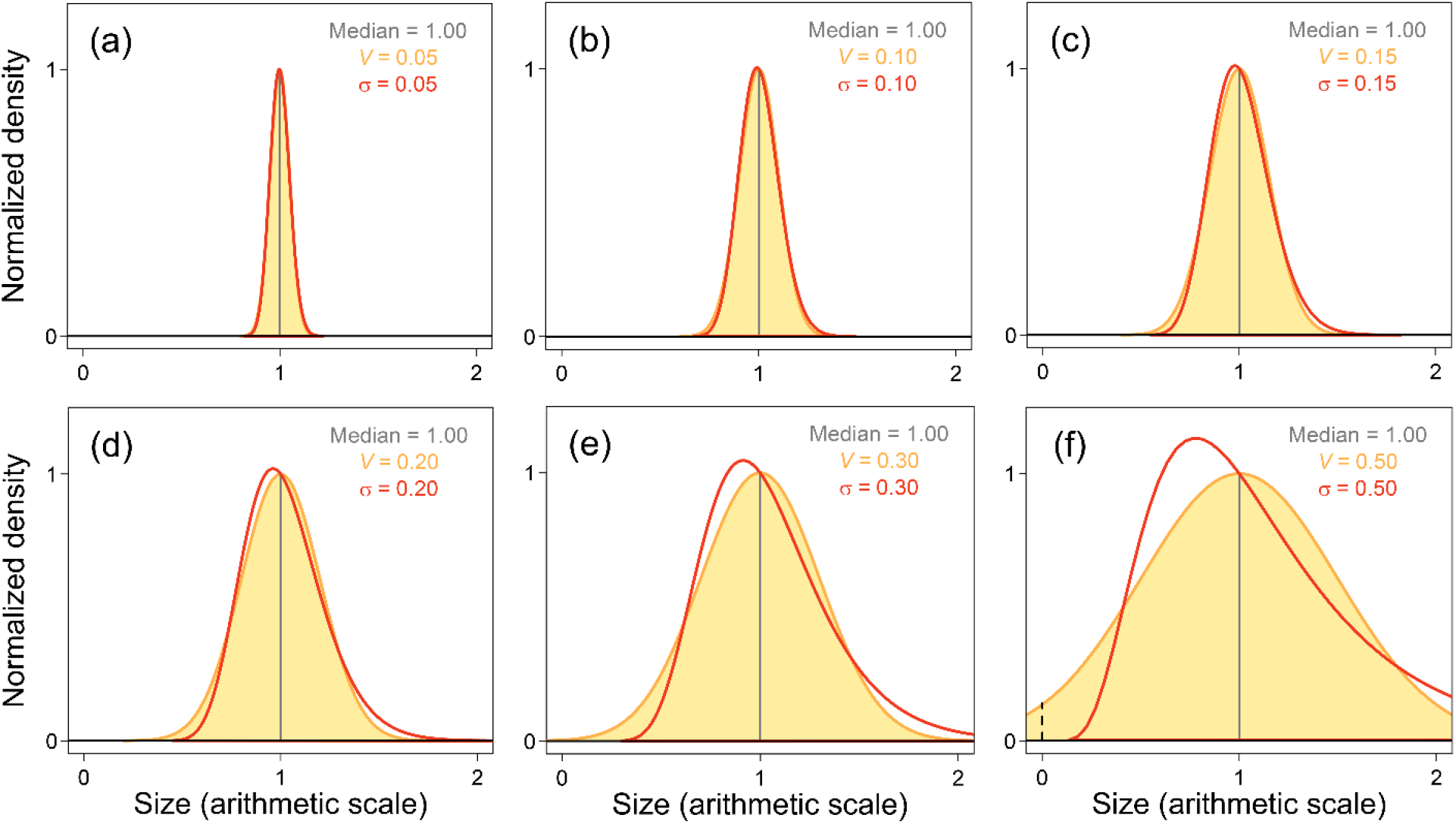
Model distributions representing biological variation on an arithmetic scale of measurement. Normal (orange) and lognormal (red) curves are superimposed for distributions with the same median size and same coefficient of variation *V* or standard deviation *σ*. Note that the two distributions are practically indistinguishable when *V* and *σ* are small (panel a) but differ substantially when *V* and *σ* are large (panels c-f). Human stature and other linear measures have *V* ≈ 0.05. Human weight and other volumetric measures have *V* ≈ 0.15. The lower tail of a distribution modeled as normal on an arithmetic scale can seemingly include impossible negative values (left of the dashed line in panel f). Empirically, biological variation is lognormal on an arithmetic scale of measurement and it becomes normal when transformed to logarithms.

### (a) Additive or multiplicative random variables

A normal distribution arises when a large number of independent and identically distributed random variables are added together or averaged, as represented by the sum *X* = *X*_1_ + *X*_2_ + … + *X*_n_. This is the central limit theorem of probability theory. A related lognormal distribution emerges when random variables are multiplied rather than added. When *X* = *X*_1_ · *X*_2_ · … · *X*_n_ we can expect *Y* = ln (*X*) to become normally distributed as the number of variables increases. This follows from the fact that *Y* = ln (*X*) = ln (*X*_1_) + ln (*X*_2_) + … + ln (*X*_n_) is itself the sum of many random variables (Otto and Day, 2007, p. 550)[8].

Whether biological variation is due to an additive combination of random variables or a multiplicative combination of random variables is an empirical question that can only be answered by study of the resulting variation. Many people, assuming that a ‘normal’ distribution of variation is canonical, have given this question little thought and assumed that additive combinations of random variables and simple normality are the rule. Others have recognized that empirical measurements yield lognormal distributions, and logarithmic transformation is required to achieve normality.

### (b) Lognormality on an arithmetic scale of measurement

Francis Galton (1879)[9] famously wrote that if you believe in an ordinary law of error, symmetrical about an arithmetic mean, then the existence of giants whose height is more than double the mean … implies the existence of dwarfs whose stature is less than nothing at all. Galton recognized that quantifying variation on a proportional logarithmic or geometric scale makes more sense than quantification on an additive arithmetic scale. The appropriate measure of location or central tendency in the proportional case is the geometric mean (exponentiated mean of the logarithms of measurements) rather than the arithmetic mean of raw measurements.

Following Galton’s lead, Cambridge mathematician Donald McAlister (1879)[10] showed that an additive law of error can be applied to the logarithms of measurements in the multiplicative geometric case just as it is commonly applied to raw measurements in the additive arithmetic case. Thus the concept of logarithmic normality was born, based on proportion rather than simple difference. A lognormal distribution is a distribution that becomes normal when its constituent elements are transformed to logarithms. Relative growth or allometry, promoted by Julian Huxley (1924)[11] and followers, is a related form of comparative analysis that becomes more tractable on logarithmic axes in recognition that underlying relationships are proportional.

John Henry Gaddum was a pharmacologist interested in the mathematics and statistics of bioassays. Observations in a bioassay vary greatly, providing insight obscured when variances are small. Gaddum (1945)[12], known for his subtle humor, started a review of lognormal distributions by observing that 18th and 19th century attempts to establish a normal law of errors were “undeservedly successful.” In Gaddum’s view the law should be a *lognormal* law of errors. He argued that incremental additions to an organism’s weight are likely to be proportional to its size. Further, when a set of objects of similar shape are measured, diameter can be normal or volume can be normal — but both cannot be normal in the same set. However, if logarithms of the diameters are normally distributed with standard deviation *s*, then logarithms of the volumes will be normal too, with standard deviation 3 · *s*. Gaddum concluded: “to convert [measurements] to logarithms before estimating their mean or variance, the usual result would be an increase in the accuracy and scope of the conclusions drawn.”

Sewall Wright (1952)[13] listed his first consideration for describing variability to be the choice of an appropriate scale — with a logarithmic scale indicated when standard deviations are proportional to means and coefficients of variation are more or less the same. For Wright, asymmetric or skewed distributions of volumes or weights also favored transformation to logarithms. Further clarity was provided by Lewontin (1966)[14], who showed that the variance of logarithms to base *e* is equivalent to the squared coefficient of variation for raw measurements. The standard deviation for the natural-log measurements is then equivalent to and a superior substitute for Pearson’s coefficient of variation. Lewontin also showed that the use of logarithms facilitates a range of statistical comparisons.

Gingerich (2000)[15] computed empirical goodness-of-fit statistics for large anthropometric samples to compare arithmetic normality and geometric normality. He found that arithmetic and geometric normality could not be distinguished for variables with small coefficients of variation. However, the two were distinguishable when coefficients of variation were larger, and, whenever distinguishable, geometric normality was consistently and strongly favored. This reinforced Galton’s and Gaddum’s shared conclusion that arithmetic measurements must be log-transformed to represent the intrinsic lognormality of biological variation appropriately.

Limpert et al. (2001)[16] summarized a review of lognormal distributions across the sciences by writing “The reasons governing frequency distributions in nature usually favor the log-normal, whereas people [seemingly] favor the normal. For small coefficients of variation, normal and log-normal distributions both fit well. For these cases, it is natural to choose the distribution found appropriate for related cases exhibiting increased variability… *This will most often be the log-normal”* (Limpert et al., 2001, p. 351, emphasis added).

Finally, Limpert and Stahel (2017)[17], echoing Galton, noted that “most quantities observed in daily life or the sciences cannot take negative values – but all normal distributions assign at least a tiny probability to this impossibility” (see, for example, the area to the left of the dashed line at zero in figure 1f). This is another reason to recognize the near ubiquity of lognormal distributions.

### (c) Logarithmic transformation

Weight distributions spanning a mouse-to-elephant spectrum of sizes are shown in figure 2. Twenty-six species are shown in figure 2a, where each has a coefficient of variation of 0.15. The same species are shown in figure 2b, where each has an equivalent standard deviation of 0.15 ln units. Each species differs from the next by a constant proportion. Species in figure 2a are lognormally distributed on an arithmetic scale of measurement. However, comparisons of species on this scale are difficult because variability changes in proportion to size, and the full spectrum of sizes and variations in relation to size are not easily illustrated on one graph. The lognormal distributions are also visibly skewed, causing separation of each sample’s mode, median, and mean.

**Figure 2.**
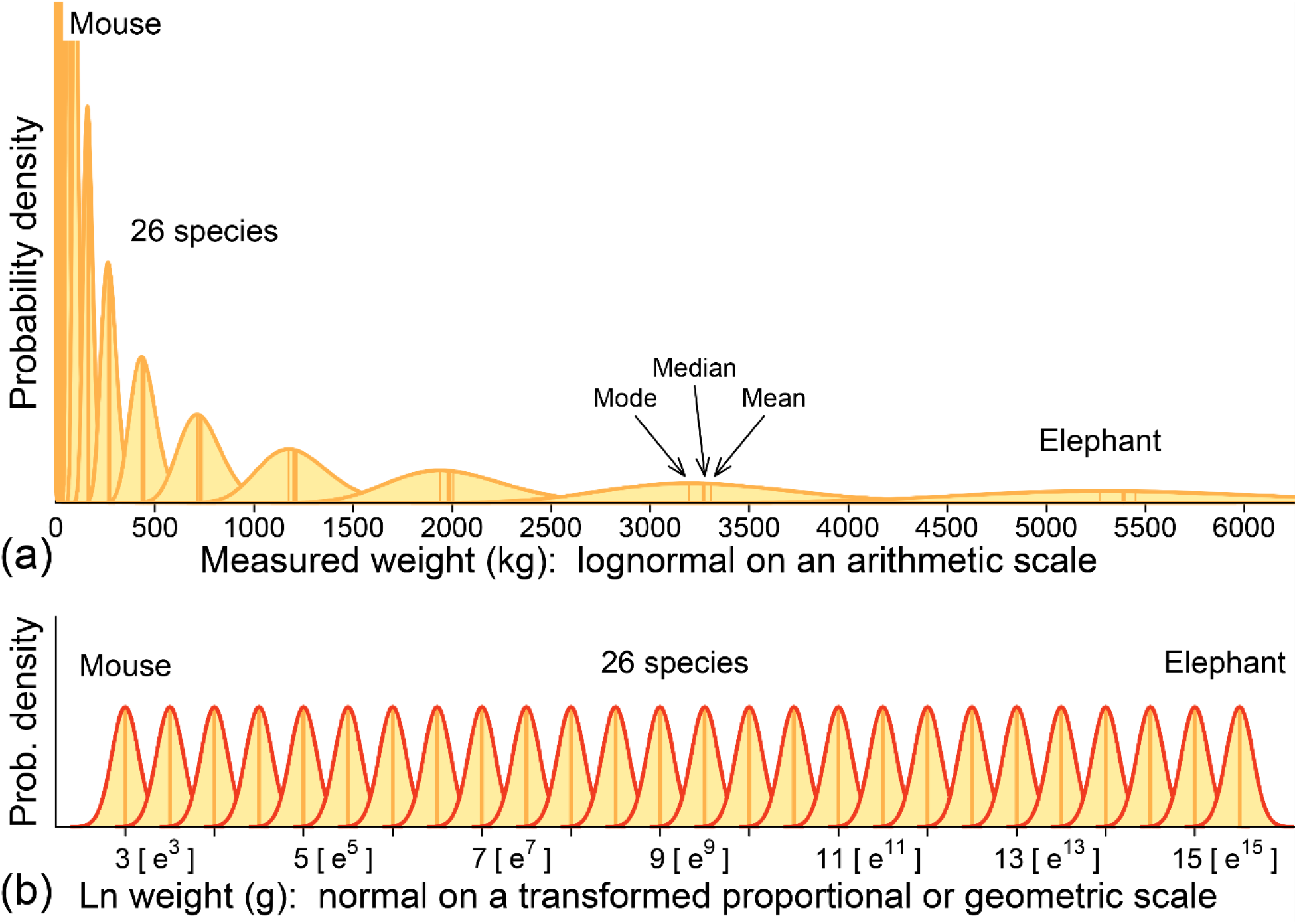
Density distributions of body weight for 26 model species spanning a mouse-to-elephant range of sizes. The distributions look very different on (a) the arithmetic scale of measurement, and (b) a proportional or geometric scale of logged measurements. Variation changes with size on an arithmetic scale (here 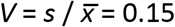) and lognormal skewness is indicated by separation of the mode, median, and mean within each sample. variation is standardized on a proportional or geometic scale (*s* = 0.15) and symmetry is indicated by coincidence of the mode, median, and mean. Note that the value 3 on a natural-log scale represents *e*^3^ on an arithmetic scale, 5 represents *e*^5^, etc., as shown in square brackets. Proportional differences are the exponentiated differences between ln values (exponentiated differences between exponents of *e*).

Species in figure 2b are shown on a proportional or geometric scale following transformation of weights to logarithms. Here comparison is easy because the distributions are normal rather than lognormal. Species have equivalent variability that is independent of size, and symmetry has replaced skewness. The mode, median, and mean values for each species now coincide.

The scale for the abscissa in figure 2b is a natural log scale, where each value corresponds to an exponent of *e*. Values ranging from 3 to 15 are shown as exponents in square brackets as a reminder of the rapidly increasing weights they represent. At the small end, 3 represents a weight of *e*^3^ = 20 grams, and at the large end 15 represents a weight of *e*^15^ = 3,269,017 grams or 3,269 kg. Differences of proportion are calculated by subtracting ln values or, equivalently, subtracting exponents of *e*. Thus *e*^3^ = 20 g and *e*^5^ = 148 g differ by a proportion of *e*^2^ = 7.389 or, when inverted, *e*^-2^ = 0.135.

Digital computers make it so easy to convert measurements to logarithms that failure to make the transformation is now inexcusable. Arithmetic lognormality and log-transformed geometric normality are equally ‘normal’ in the sense that both fit our measurements, but the standardized variability and symmetry of geometric normality greatly simplifies analysis.

## 3. Zero Force Evolutionary Law on a Proportional Scale

McShea et al. (2019) formulated their zero-force expectation of evolutionary divergence in terms of additive change on an arithmetic scale, but acknowledged that change in biology is often proportional and appropriately interpreted on a logarithmic scale. Authors cited in the previous section on arithmetic lognormality and geometric normality would say that a geometric or logarithmic scale is not optional: evolutionary change is always proportional and can only be interpreted on a logarithmic scale. Three paired random-walk simulations illustrate the statistical expectation that initial difference will be maintained in each case, demonstrating that no systematic divergence is expected between randomly evolving lineages.

### (a) Random walks with no initial difference

The simulation in figure 3 shows two 100-step random walks that are realistically scaled from a biological point of view. Lineage A in yellow and lineage B in orange both start at size *μ_A_* = 2.5 on a natural-log proportional scale, with standard deviation *σ* = 0.1 and variance *σ*^2^ = 0.01 (red normal curve at generation zero). Note that *σ* = 0.1 is equivalent to a coefficient of variation *V* = 0.10 or 10% on an arithmetic scale. Change at each step, positive or negative, averages 0.08 standard deviations per generation, which is conservative rate of change (Gingerich 2019)[18]. After 100 generations lineage A is at *x_A_* = 1.265, which is well to the left of the starting size (dashed red line) and proportionally smaller than expectation after 100 generations (represented by the broad orange normal curve at the top of figure 3a centered at *x_A_* = 2.5 with standard deviation *s* = 1 and variance *s*^2^ = 1).

**Figure 3.**
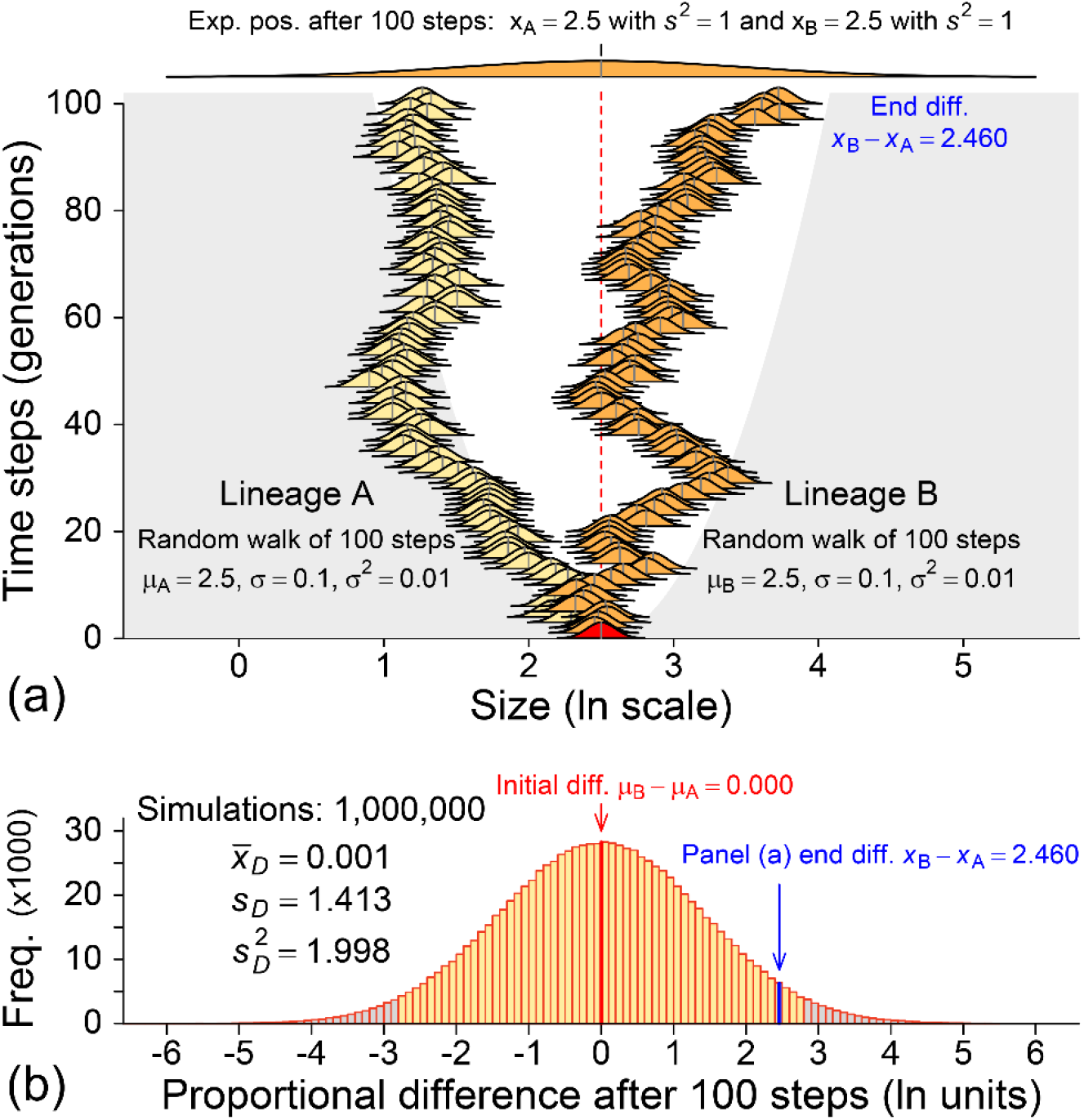
Accumulation of proportional difference for random walks that start with no difference. (a) Simulation shows two random walks. Lineage A (yellow) and lineage B (orange) both start at size *μ_A_* = *μ_B_* = 2.5 (‘size’ can be linear, areal, or volumetric in any appropriate unit). The initial difference is *μ_B_* – *μ_A_* = 0. The end difference at step 100 is *x_B_* – *x_A_* = 3.725 – 1.265 = 2.460 (or *e*^3-725^ / *e*^1.265^ = *e*^2.460^), which means B ends 11.705 times the size of A. The expected value for a random walk is the initial value (dashed line), but realized values can vary greatly. The total variance in each lineage increases in proportion to the number of steps: here from *σ*^2^ = 0.01 to *σ*^2^ = 1 (shown by the broad orange normal curve at the top of panel a. (b) Histogram of proportional differences for a million random-walk simulations like the one in panel a. The mean variance for the simulations, *s_D_*^2^ = 1.998, is almost exactly the sum of variances for each pair of 100-step random walks. The mean difference, 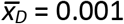, is almost exactly the initial difference *μ_B_* – *μ_A_* = 0.000. Gray shading encloses a 95% confidence interval in each panel. Negative differences balance positive differences, and there is no statistical expectation of divergence for lineages evolving randomly thorough time.

After 100 generations lineage B is at *x_B_* = 3.725, which is well to the right of expectation when compared to the orange normal curve at the top of the figure. The proportional difference between *x_A_* and *x_B_* after 100 generations is *x_B_* – *x_A_* = 3.725 – 1.265 = 2.460. Exponentiating, this means that B at generation 100 is *e*^2.460^ = 11.705 times the size of A. Inverting the comparison, A at generation 100 is *e*^-2.460^ = 0.085 times the size of B. We can put this proportional difference of 2.460 in context by running the random-walk simulation many times.

The histogram in figure 3b, normally distributed, summarizes results from a million iterations of the random-walk simulation of figure 3a. The average difference, 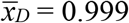, for all the simulations is almost exactly the initial difference *μ_B_* – *μ_A_* = 0.000, and the pooled variance for all the simulations, *s_D_*^2^ = 1.998, is almost exactly the sum of the variances at each step in the two lineages combined (2 · 100 · 0.01 = 2.000). A proportional difference of *x_B_* – *x_A_* = 2.460 is large, but it is less than two standard deviations from the mean 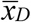 for the simulated differences.

Random-walk lineages like those in figure 3a will always diverge to some degree, but the divergence of one with respect to the other can be positive or negative and the most likely divergences are the smallest. This is why the expected difference *x_B_* – *x_A_* between the lineages in any generation remains the initial difference *μ_B_* – *μ_A_*. In figure 3b the expected difference *x_B_* – *x_A_* = *μ_B_* – *μ_A_* = 0.000.

### (b) Random walks with an initial difference and no crossing

The simulation in figure 4 shows two 100-step random walks like those in figure 3. In figure 4a, modeled after figure 5 of McShea et al. (2019), lineage A in yellow starts at size *μ_A_* = 2.0 and lineage B in orange starts at size *μ_A_* = 3.0 on a natural-log proportional scale (starting distributions are shown in red at generation zero). Both have standard deviation *σ* = 0.1 and variance *σ*^2^ = 0.01. Change at each step, positive or negative, averages 0.08 standard deviations per generation. After 100 generations lineage A is at *x_A_* = 1.382, which is a little to the left of the starting size (dashed red line), and proportionally a little smaller than expectation after 100 generations (represented by the broad yellow normal curve at the top of figure 4a centered at *x_A_* = 2.0 with standard deviation *s* = 1 and variance *s*^2^ = 1).

**Figure 4.**
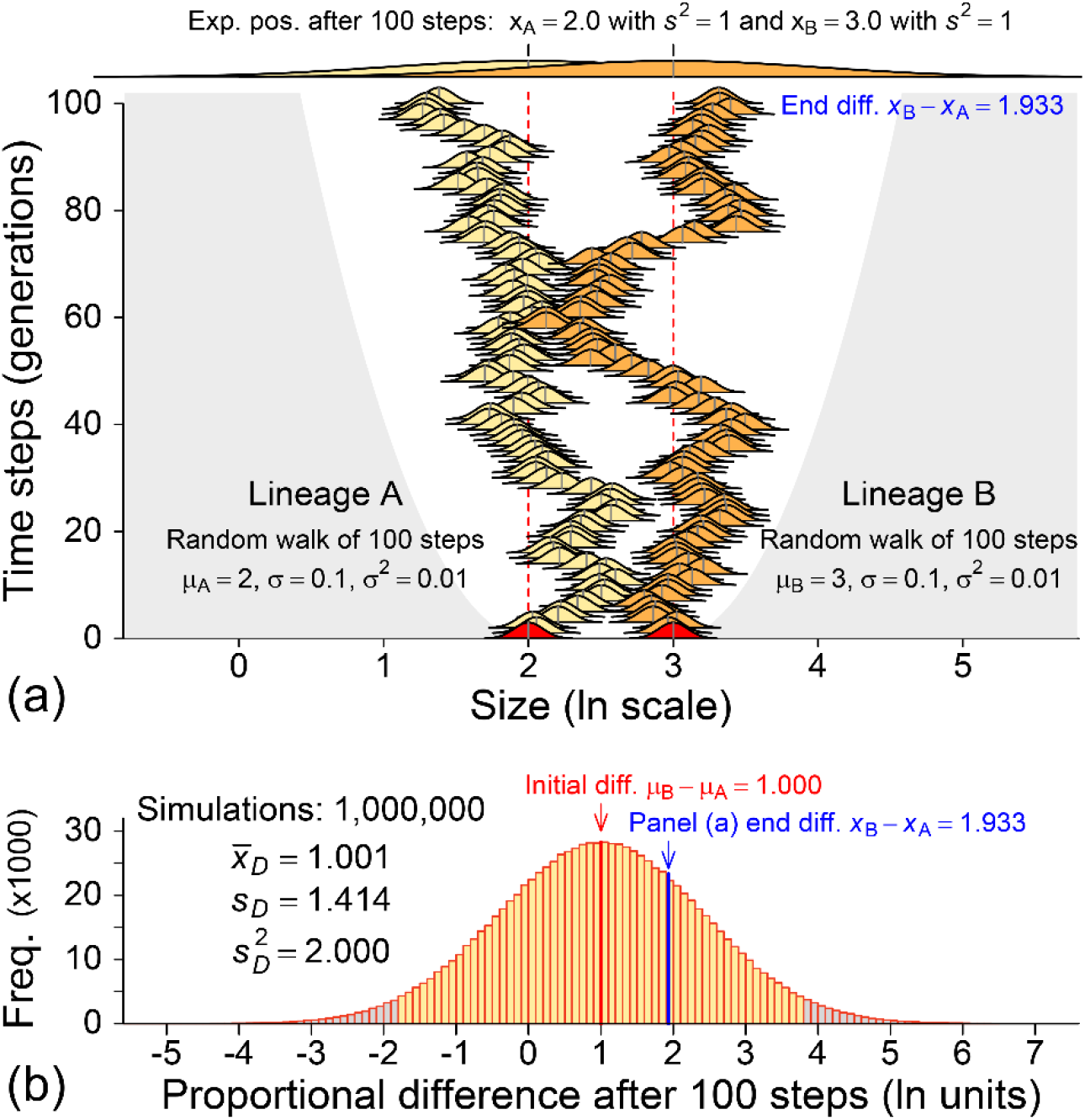
Accumulation of proportional difference for random walks that start with a difference. (a) Simulation shows two random walks that do not cross. Lineage A (yellow) and lineage B (orange) start at size *μ_A_* = 2 and size *μ_B_* = 3, respectively (‘size’ can be linear, areal, or volumetric in any appropriate unit). The initial difference is *μ_B_* – *μ_A_* = 3 – 2 = 1 (or *e*^3^ / *e*^2^ = *e*^1^), which means B starts 2.718 times the size of A. The end difference at step 100 is *xB – x_A_* = 3.315 – 1.382 = 1.933 (or *e*^3-315^ / *e*^1-382^ = *e*^1-933^), which means B ends 6.910 times the size of A. The expected value for each random walk is its initial value (dashed line), but realized values can vary greatly. The total variance in each lineage increases in proportion to the number of steps: here from *σ*^2^ = 0.01 to *σ*^2^ = 1 (shown by the broad overlapping yellow and orange normal curves at the top of the panel). (b) Histogram of proportional differences for a million random-walk simulations like the one in panel a. The mean variance for the simulations, *s_D_*^2^ = 2.000, is the sum of 100-step variances for each pair of random walks. The mean proportional difference, 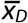 = 1.001, is almost exactly the initial difference *μ_B_* – *μ_A_* = 1.000. Gray shading encloses a 95% confidence interval in each panel. Negative differences balance positive differences, and there is no statistical expectation of divergence for lineages evolving randomly thorough time.

After 100 generations lineage B is at *x_B_* = 3.315, which is a little to the right of expectation when compared to the orange normal curve at the top of the figure. The proportional difference between *x_A_* and *x_B_* after 100 generations is *x_B_* – *x_A_* = 3.315 – 1.382 = 1.933. Exponentiating, this means that B at generation 100 is *e*^1.933^ = 6.910 times the size of A. Inverting the comparison, A at generation 100 is *e*^-1.933^ = 0.145 times the size of B. Here again, we can put a proportional difference of 1.933 in context by running the random-walk simulation many times.

The histogram in figure 4b summarizes results from a million iterations of the randomwalk simulation of figure 4a. The average difference, 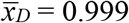, for all the simulations is almost exactly the initial difference *μ_B_* – *μ_A_* = 1.000, and the pooled variance for all the simulations, *s_D_*^2^ = 2.002, is almost exactly the sum of the variances at each step in the two lineages combined (2 · 100 · 0.01 = 2.000). A proportional difference of *x_B_* – *x_A_* = 1.933 is large, but it is less than one standard deviation from the mean 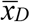 for the simulated differences.

Random-walk lineages like those in figure 4a can converge or diverge. The changes of one with respect to the other can be positive or negative and the most likely changes are the smallest. This is why the expected difference *x_B_* – *x_A_* between the lineages in any generation remains the initial difference *μ_B_* – *μ_A_*. In figure 4b the expected difference *x_B_* – *x_A_* = *μ_B_* – *μ_A_* = 1.000.

### (c) Random walks with an initial difference and crossing

The simulation in figure 5 shows two 100-step random walks like those in figures 3 and 4. In figure 5a, modeled after figure 5 of McShea et al. (2019), lineage A in yellow starts at size *μ_A_* = 2.0 and lineage B in orange starts at size *μ_A_* = 3.0 on a natural-log proportional scale (starting distributions are shown in red at generation zero). Both have standard deviation *σ* = 0.1 and variance *σ*^2^ = 0.01. Change at each step, positive or negative, averages 0.08 standard deviations per generation. After 100 generations lineage A is at *x_A_* = 3.023, which is well to the right of the starting size (left dashed red line), and proportionally larger than expectation after 100 generations (represented by the broad yellow normal curve at the top of figure 5a centered at *x_A_* = 2.0 with standard deviation *s* = 1 and variance *s*^2^ = 1).

**Figure 5.**
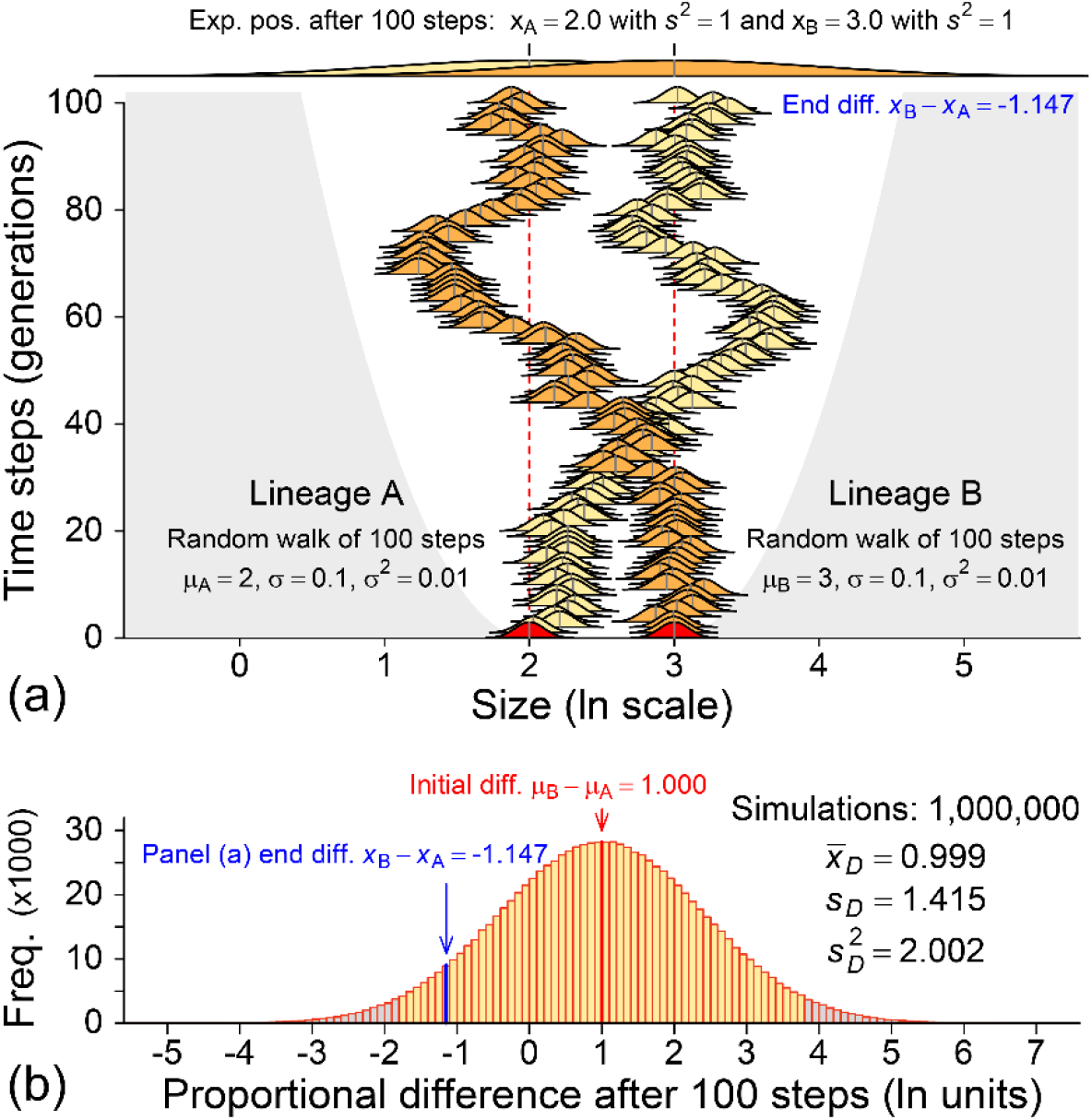
Accumulation of proportional difference for random walks that start with a difference. (a) Simulation shows two random walks that cross. Lineage A (yellow) and lineage B (orange), start at size *μ_A_* = 2 and size *μ_B_* = 3, respectively (‘size’ can be linear, areal, or volumetric in any appropriate unit). The initial difference is *μ_B_* – *μ_A_* = 3 – 2 = 1 (or *e*^3^ / *e*^2^ = *e*^1^), which means B starts 2.718 times the size of A. The end difference at step 100 is *x_B_* – *x_A_* = 1.875 – 3.023 = −1.147 (or *e*^1.875^ / *e*^3.023^ = *e*^-1.147^), which means B ends 0.317 times the size of A. The expected value for each random walk is its initial value (dashed line), but realized values can vary greatly. The total variance in each lineage increases in proportion to the number of steps: here from *σ*^2^ = 0.01 to *σ*^2^ = 1 (shown by the broad overlapping yellow and orange normal curves at the top of the panel). (b) Histogram of proportional differences for a million random-walk simulations like the one in panel a. The mean variance for the simulations, *s_D_*^2^ = 2.002, is almost exactly the sum of 100-step variances for each pair of random walks. The mean proportional difference, 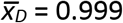, is almost exactly the initial difference *μ_B_* – *μ_A_* = 1.000. Gray shading encloses a 95% confidence interval in each panel. Negative differences balance positive differences, and there is no statistical expectation of divergence for lineages evolving randomly thorough time.

After 100 generations lineage B is at *x_B_* = 1.875, which is well to the left of expectation when compared to the orange normal curve at the top of the figure. The proportional difference between *x_A_* and *x_B_* after 100 generations is *x_B_* – *x_A_* = 1.875 – 3.023 = −1.147. Exponentiating, this means that B at generation 100 is *e*^-1.147^ = 0.317 times the size of A. Inverting the comparison, A at generation 100 is *e*^1.147^ = 3.152 times the size of B. We can put a proportional difference of −1.147 in context by running the random-walk simulation many times.

The histogram in figure 5b, a normal distribution, summarizes results from a million iterations of the random-walk simulation of figure 5a. The average difference, 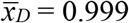, for all the simulations is almost exactly the initial difference *μ_B_* – *μ_A_* = 1.000, and the pooled variance for all the simulations, *s_D_*^2^ = 2.002, is almost exactly the sum of the variances at each step in the two lineages combined (2 · 100 · 0.01 = 2.000). A proportional difference of *x_B_* – *x_A_* = −1.147 is small, but it is less than two standard deviations from the mean 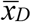 for the simulated differences.

Random-walk lineages like those in figure 5a can converge or diverge. The changes of one with respect to the other can be positive or negative and the most likely changes are the smallest. This is why the expected difference *x_B_* – *x_A_* between the lineages in any generation remains the initial difference *μ*B – *μ*A. In figure 5b the expected difference *x_B_* – *x_A_* = *μ_B_* – *μ_A_* = 1.000.

Additional simulations could be carried out, with shorter or longer lineages, smaller or larger rates of change at each step, and lesser or greater initial differences between lineages, but the overall result would be the same. On a scale of proportional difference, the expected difference at the end of any simulation run, which is the mean difference 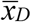 for a large number of runs, will always be the initial difference. There is no expectation of a systematic zero-force increase in proportional difference for random walks evolving thorough time.

## 4. Discussion

Interpretation often depends on the choice of scale or reference frame for a particular analysis, and the scale must be appropriate. If phenotypic variation and evolution involved additive combinations of random variables, then it might be appropriate to study normality and frame a zero-force evolutionary law on an arithmetic scale of measured amounts. However, as reviewed above, phenotypic variation is found, empirically, to be lognormally distributed when plotted on the arithmetic scale of measurement (Galton 1879; Gaddum 1945; Wright 1952; Gingerich 2000; Limpert et al. 2001; Limpert and Stahel 2017)[9, 12, 13, 15–17]. Thus the underlying combinations of random variables are multiplicative rather than additive, and normality requires that measurements be converted to logarithms before plotting on what is then a proportional or geometric scale. Analysis on an additive arithmetic scale might be ‘good enough’ when variability is low (as in figure 1a), but it makes no sense to generalize based on this ambiguous indeterminacy.

Phenotypic variation is proportional to mean size, and a normally or lognormally distributed population may approach zero in size but never cross it to become negative. Figure 2a illustrates this. There is no negative measured size. It may be less obvious, but the value of a difference between populations is similarly proportional to size. Differences can approach zero, but never cross zero to become negative.

McShea et al. (2019) chose an arithmetic scale and differences of amount to represent evolutionary change and to frame their zero-force evolutionary law. They scaled variability as if it is normal on an arithmetic scale, ignoring its proportionality to mean size and in the process ignoring the proportionality of differences to mean size. McShea et al. recognized that crossover differences on an arithmetic scale represent distances that should not be negative. Their solution was to reflect the negative differences as absolute values, and then add these to the corresponding distribution of positive differences. This reflection is the source of their zero-force expectation of increasing divergence separating lineages through time. Other people, starting with Galton (1879), have recognized that negative differences render an arithmetic scale untenable, and then reformulated their analyses on a proportional scale.

Randomly evolving lineages are most easily and appropriately modeled as normally distributed on a proportional geometric or logarithmic scale. When these diverge after starting at the same initial size, as in figure 3, fully half of the divergences will involve a negative difference *x_B_* – *x_A_* < 0. When randomly evolving lineages converge or diverge from different initial sizes, as in figures 4 and 5, some crossover of large and small lineages is expected, which can also yield *x_B_* – *x_A_* < 0. The important point is that any acquisition of differences less than the initial difference between lineages will tend to be balanced by a complementary acquisition of differences that are greater than the initial difference.

We can go a step farther and calculate confidence limits for the proportional differences between lineages after *N* steps or generations, based on the initial means *μ_A_* and *μ_B_* and their common variance *σ*^2^. The expected difference 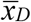 after *N* steps is 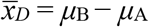, the initial difference, and the expected variance *s_D_*^2^ after *N* steps is *s_D_*^2^ = 2 · *N* · *σ*^2^. A 95% confidence interval for the proportional difference at any step *N* is given by 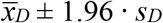, which ranges from −2.772 to +2.772 for *N* = 100 in figure 3b and from −1.772 to +3.772 for *N* = 100 in figures 4b and 5b. These 95% confidence intervals fall to −2.772 ln units below the expected difference and rise to 2.772 ln units above the expected difference. Exponentiating, each *N* = 100 generation confidence interval ranges from a factor of about 1/16 below expectation to a factor of about 16 above expectation. The expected proportional difference between lineages remains the initial difference, but the corresponding confidence interval expands rapidly in proportion to the square-root of *N*.

Based on these results we can restate the zero-force evolutionary law as a null model. Figures 3–5, scaled proportionally, illustrate this. Given two entities evolving randomly and independently — in the absence of forces or constraints acting on the lineages or on differences between them — the statistical expectation is that there will be no net change in the distance between the lineages, and there is a corresponding statistical expectation of no systematic divergence of lineages.

